# Branching architecture affects genetic diversity within an individual tree

**DOI:** 10.1101/2023.10.02.560431

**Authors:** Sou Tomimoto, Yoh Iwasa, Akiko Satake

## Abstract

**Premise:** While a tree grows over many years, somatic mutations accumulate and generate a genetically diverse individual. Trees can transmit such mutations to subsequent generations, potentially enhancing the genetic diversity of the population. We study a mathematical model to understand the relationship between within-individual genetic variation and branching architecture.

**Methods:** We generate branching architecture by repeatedly adding two new branches (main and lateral daughter branches) to each terminal branch (mother branch). Tree shape is determined by two key parameters: daughter-mother ratio (DM) and main-lateral ratio (ML). During branch elongation, somatic mutations accumulate in the stem cells of a shoot apical meristem (SAM) at the tip of each branch. In branching, all the stem cells are passed on from the mother to the main daughter branch, but only one stem cell is chosen for the lateral daughter branch. We evaluate genetic variation by 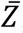, the mean genetic differences between all pairs of branches of a tree, and examine how 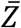 varies with DM and ML while keeping the total branch length constant.

**Key results and Conclusions:** (1) 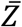 attains the maximum for an intermediate DM, when stem cells in the SAM are genetically homogeneous; (2) 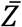 decreases monotonically with DM when stem cells are genetically heterogeneous; and (3) 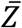 increases monotonically with ML. Even though the total branch length remains constant, the within-individual genetic variation differs substantially. The results demonstrate the importance of branching architecture in storing genetic diversity.

## INTRODUCTION

A long-lived tree accumulates somatic mutations during its growth, leading to genetic variation among its branches. Because trees lack a clear distinction between germline and soma, somatic mutations can be transmitted to subsequent generations (Whitham and Slobodchikoff, 1981; Sutherland and Watkins, 1986; Plomion et al., 2018) and contribute to the genetic diversity in the tree population (Antolin and Strobeck, 1985; Schoen and Schultz, 2019).

Recent advances in genomic technology, such as next-generation sequencing, have facilitated the quantitative identification of somatic mutations in long-lived modular organisms like trees (Reusch et al., 2021). The presence of genetic variation within an individual has been reported in numerous tree species (Schmid-Siegert et al., 2017; Plomion et al., 2018; Wang et al., 2019; Hanlon et al., 2019; Orr et al., 2020; Hofmeister et al., 2020; Perez-Roman et al., 2021; Duan et al., 2022; Satake et al., 2023). Somatic mutations accumulated as trees age (Satake et al. 2023) and were distributed following the branching architecture of a tree (Schmid-Siegert et al., 2017), and the phylogeny of somatic mutations in a single tree showed the same topology as the physical shape of a tree (Perez-Roman et al., 2021; Satake et al. 2023). Genetic variation seemingly presented not only among branches but also among cells in a shoot apical meristem (hereafter SAM) of a single branch (Zahrdadníková et al., 2020; Duan et al., 2022; Schmitt et al., 2022). Additionally, somatic mutations were mostly neutral within a tree (Orr et al., 2020; Perez-Roman et al., 2021; Duan et al., 2022; Satake et al., 2023). These mutations were transmitted to offspring (Plomion et al., 2018), and at the stage of seed production and seedling growth, they seem to have experienced natural selection (Ally et al., 2016; Satake et al., 2023). These empirical studies have significantly expanded our understanding of how raw materials of evolution, novel mutations, emerge and proliferate in naturally growing trees. However, the influence of the architecture of a tree on somatic mutation accumulation remains unresolved.

The branching architecture of trees exhibits remarkable diversity depending not only on species and genotypes but also on local environments that individuals have undergone during growth (Fisher and Hibbs, 1982; Sussex and Kerk, 2001; Barthélémy and Caraglio, 2007). The branching architecture may significantly influence the pattern of mutation accumulation and the amount of genetic variation formed within an individual tree, because mutations occur and accumulate while a tree shapes its architecture through shoot elongation and branching. These growing processes involve complex dynamics of stem cells within a SAM and somatic selection on mutations. To understand such complex cellular-level dynamics, mathematical models proved valuable. Several mathematical models have unveiled the impact of various factors on somatic mutation accumulation, such as the cellular structure of a SAM (Klekowski and Kazarinova-Fukshansky, 1984a; Klekowski et al, 1985; Pineda-Krch and Lehtilä, 2002; Iwasa et al. 2023a), tree ontogeny (Klekowski et al., 1989; Tomimoto and Satake, 2023), and somatic selection on mutated stem cells and branches (Klekowski and Kazarinova-Fukshansky, 1984b; Antolin and Strobeck, 1985; Pineda-Krch and Fagerström, 1999; Otto and Orive, 1995; Folse and Roughgarden, 2012). However, it remains unclear how branching architecture influences (1) the accumulation pattern of somatic mutations and (2) the amount of genetic variability stored within an individual tree.

In this paper, we studied a simple mathematical model to address these questions. We generated different branching architectures using a bifurcation tree and quantified the genetic difference between pairs of SAMs among branches in a single tree. In addition to branching architectures, our model highlighted the importance of the genetic heterogeneity within the stem cell population in a SAM, which results from the dynamics of stem cells during shoot elongation and branching. For a genetically homogeneous SAM, we found that the genetic difference between two branches is simply proportional to the length of a path connecting the SAM of these two branches. For a genetically heterogeneous SAM, genetic differences become larger than a homogeneous SAM, with their degree depending on a branching architecture. We then evaluated the within-individual genetic variation by the mean of the genetic differences over all branch pairs and addressed how this value changes under different branching architectures. Our results demonstrated how branching architecture affects the genetic structure within a tree depending strongly on the genetic heterogeneity within a SAM.

## MATERIALS AND METHODS

### Branching architecture

To model branching architecture, we adopt a two-dimensional bifurcation tree simplifying the model developed in previous studies (Honda, 1971; Honda and Fisher, 1978; Honda and Fisher, 1979; Fisher and Honda, 1979). We model one step of bifurcation in the following events: (1) A mother branch produces two daughter branches, main and lateral ones. (2) The lengths of the main and lateral branches are given in the ratios *r*_1_ and *r*_2_ to the mother branch, respectively. (3) Whereas the main branch grows straight from the mother branch, the lateral branch grows with angle *θ* (Fig. 1A).

**Figure 1.**
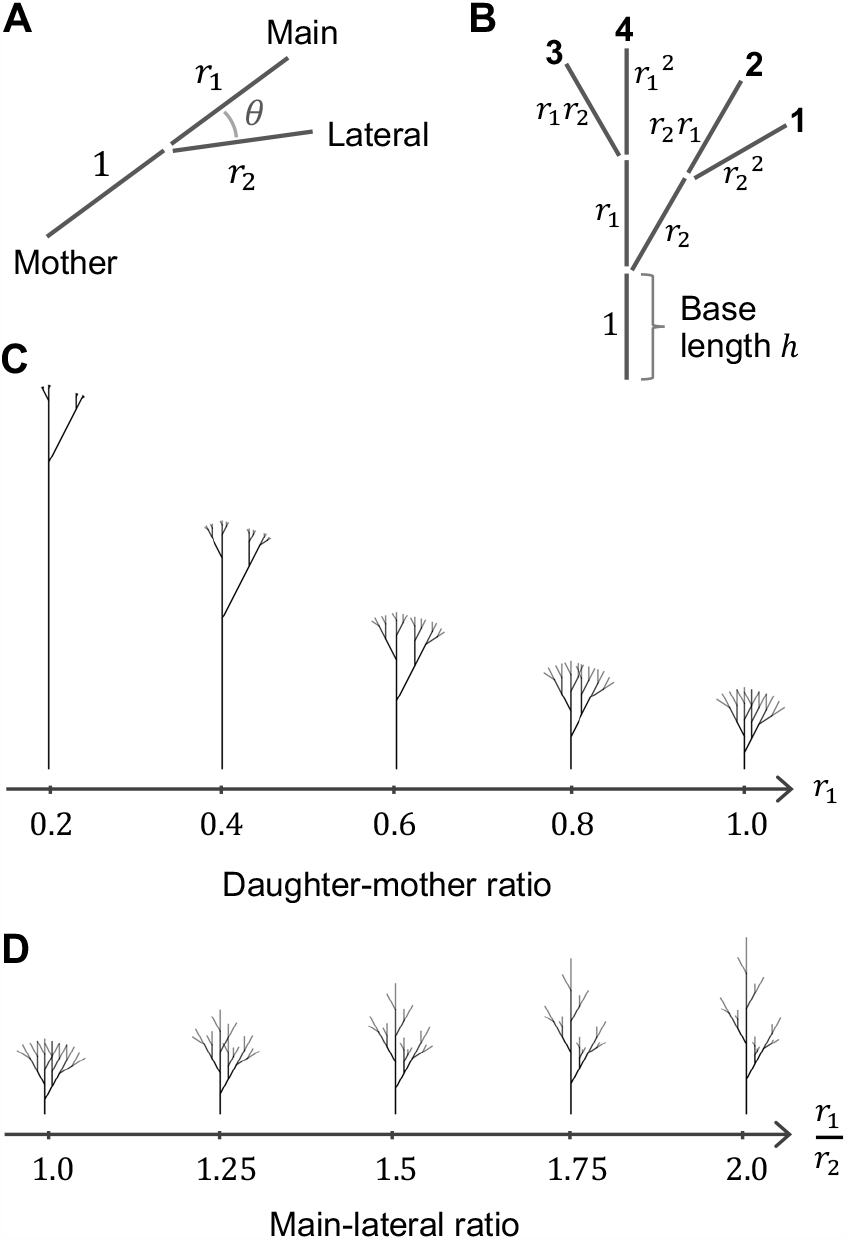
Branching architecture. (A) Illustration of bifurcation. A mother branch produces main and lateral branches whose lengths are in the ratio *r*_1_ and *r*_2_ to the mother branch, respectively. The main branch straightly elongates from the mother branch, whereas the lateral branch elongates with angles *θ*; (B) Illustration of the branching architecture formed by two reiterations of the bifurcation process (*N* = 2). Bold numbers (1–4) denote branch IDs. The total length of branches is kept constant by adjusting the base length h for different branching architectures. The relative length of branches to the base length is noted next to each branch; (C) Branching architectures varying the daughter-mother ratio, under *r*_1_/*r*_2_ = 1; (D) Branching architectures varying the main-lateral ratio, under *r*_1_ = 1. Other parameters were *N* = 4 and *θ* = 30°. See Honda (1971) for graphing.

Reiteration of the bifurcation step shapes the branching architecture. In one step of bifurcation, each mother branch at the terminal portion of a tree produces two daughter branches. Then, the mother becomes a nonterminal branch and the daughters become the two terminal branches, which subsequently produce their own daughter branches. The number of terminal branches doubles in each reiteration step. After *N* times bifurcation, a single branch thus produces 2^*N*^ terminal branches (Fig. 1B).

### (1) Daughter to mother ratio and main to lateral ratio

For different branching architectures, we focus on changes in the length of a branch unit (an elongated portion of a branch between bifurcation events) and control branch length using two quantities that determine the architecture: the ratio of daughter to mother branches, *r*_1_, and the ratio of main to lateral daughter branches, *r*_1_/*r*_2_. These quantities sufficiently describe diverse branching architectures.

The daughter-mother ratio *r*_1_, or DM ratio for short, reflects the relative length of lateral branches. After the *N* times reiteration, branches at the edge of the tree are 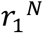 times shorter than the length of the base trunk (first mother branch). As the DM ratio *r*_1_ decreases, the branching architecture changes its crown shape from dense to sparse and its height from short to tall. Additionally, the length of the trunk increases (Fig. 1C). Here, we examine the model with *r*_1_ < 1. The main-lateral ratio *r*_1_/*r*_2_, or ML ratio, determines the degree of apical dominance, or apical control (Cline, 1997). The high ratio indicates strong apical dominance, with the lateral branch much shorter than a main branch. As the ML ratio increases, the branching architecture changes its crown shape from round to conic (Fig. 1D). Here, we examine the model with *r*_1_/*r*_2_ > 1.

### (2) Total length of branches in an individual tree

In comparing the different branching architectures, the total length of branches is kept constant by adjusting the base length h, the branch length from the base to the first branching event (Fig. 1B). This implies that the expected number of total mutations in a whole tree is the same between different branching architectures, because mutations are assumed to occur at a rate proportional to a physical branch length (see discussion below). We adjust h as follows:

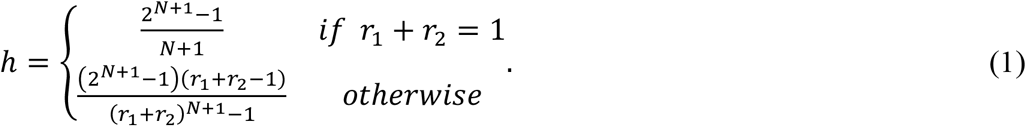

By choosing this way, the total length of branches becomes independent of both *r*_1_ and *r*_2_, and is given by 2^*N*+1^ − 1 (See Appendix A in the Supplemental Material). We calculated the within-individual genetic variation for branching architectures that differ in *r*_1_ and *r*_1_/*r*_2_ (Fig. 1B, C, D). Notably, we focus on the length of branches, because the accumulation of mutations requires the elongation of branches in the model. The branching angle *θ* is thus irrelevant in our analysis.

### Genetic structure within a tree

#### (1) Tree growth and accumulation of somatic mutations

A shoot apical meristem (SAM) positions at the tip of each branch and includes a small number of initial cells, or stem cells (Dermen, 1969). Each stem cell undergoes asymmetric cell division and produces one successor stem cell and a differentiated cell, which proliferates and leads to shoot elongation (Stewart and Dermen, 1970; Umeda et al., 2021). Any portion of the shoot below the tip consists of cells that originated from the SAM sometime before. Consequently, the physical location of the shoot (lower to upper) corresponds to the time for the asymmetric cell division of the stem cell (earlier to later) (Iwasa et al., 2023a). As a shoot grows, stem cells in the SAM involve somatic mutations. The cells in SAMs at the tips of a tree thus accumulate more mutations than cells in the tree base, and different branches independently accumulate mutations. Mutations occurring in these stem cells shape genetic structure within a tree and can be inherited to offspring. Our model thus focuses on somatic mutations that accumulate in these stem cells in SAMs.

The growth of trees involves two processes: elongation and branching (Antolin and Strobeck, 1985; Tomimoto and Satake, 2023). In elongation, apical growth occurs through repeating cell divisions in the SAM, during which somatic mutations occur and accumulate. In branching, a new lateral daughter branch is formed from an old main branch. In this process, the meristem of a lateral branch, called axillary meristem, is formed by stem cells in the apical meristem of the main branch (Poething, 1989; Sussex and Kerk, 2001). Thus, somatic mutations that accumulate in the apical meristem are stochastically sampled and passed on to the axillary meristem, depending on the sampled stem cells that contribute to the formation (Burian et al., 2016; Tomimoto and Satake, 2023).

#### (2) Mutation accumulation during elongation

During elongation, somatic mutations occur and accumulate in each stem cell lineage. For simplicity of the argument, we first consider the process of mutation accumulation in a single stem cell lineage. Let *d* represent the distance of branch elongation, the expected number of accumulated mutations is given as follows:

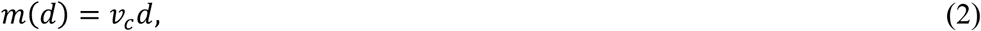

where *v*_+_ is the mutation rate per cell (whole genome) per unit distance branch elongation (for derivation see Appendix B in the Supplemental Material). We note this rate accounts for both cell-division and time-dependent mutations, assuming uniform time requirement for achieving unit growth across all branches.

To extend the model of mutation accumulation in a single cell lineage to a SAM having multiple stem cells, we need to consider the dynamics of stem cell lineages. Previous studies have considered two contrasting dynamics of stem cells during elongation: stochastic and structured meristems that lead to genetically homogenous and heterogenous SAM, respectively (Klekowski and Kazarinova-Fukshansky, 1984a; Tomimoto and Satake, 2023). If the SAM has a small number of stem cells and one stem cell lineage frequently replaces another lineage during elongation, a single cell lineage quickly becomes fixed in the SAM due to somatic genetic drift (Klekowski and Kazarinova-Fukshansky, 1984a; Iwasa et al., 2023a). In such stochastic meristem, only mutations that occur in this single lineage accumulate in the SAM, resulting in genetic homogeneity (Tomimoto and Satake, 2023). In contrast, if stem cells engage in asymmetric cell divisions and stem cell lineages are kept separate, somatic mutations independently accumulate in each cell lineage, leading to high genetic heterogeneity within the SAM (Tomimoto and Satake, 2023; Iwasa et al., 2023b). As an extreme case of this structured meristem, our model assumes that (i) all stem cell lineages are retained during elongation. Hence, stem cells in the SAM at the trunk tip have their common ancestor only at the base of the tree. Whether a tree has a stochastic or structured meristem may depend on species (Tomimoto and Satake, 2023). In the present paper, we represent the stochastic meristem with homogeneity as the case with a single stem cell lineage (*α* = 1), and the structured meristem with heterogeneity as the case with multiple stem cell lineages (*α* > 1).

#### (3) Sampling of a cell in branching

In branching, the axillary meristem of a new lateral branch is formed by some stem cells in the apical meristem of the main branch (Poething, 1989; Sussex and Kerk, 2001). Following Tomimoto and Satake (2023), we make two assumptions: (ii) The apical meristem retains all stem cell lineages during branching (i.e., monopodial branching). (iii) A newly formed axillary meristem originates from a single stem cell sampled from the apical meristem. Hence, only mutations accumulated in this sampled cell lineage are passed to and fixed in the axillary meristem losing genetic heterogeneity. In contrast, the apical meristem retains all stem cell lineages and the genetic heterogeneity that have accumulated during prior elongation.

#### (4) Index of genetic distance and variation within a tree

As an index for evaluating the accumulation pattern of somatic mutations within an individual tree, we first consider the number of mutations that differ between SAMs at two terminal branches. We define the “pairwise genetic difference” as the average heterozygosity between cells randomly sampled from the SAMs of each branch. The fraction of mutated stem cells within the meristem is represented as 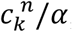, where *α* is the number of stem cells and 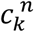 is the number cells that mutated at site *k* in the SAM of branch *n*. The probability that a mutated stem cell is sampled from only one of the two branches, *n* and *m*, can be given as 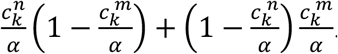. By summing this quantity across all sites of the genome, we obtain the difference between branches *n* and *m, Z*_*n,m*_, as follows:

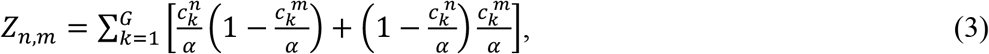

where *G* represents the genome size. Notably, *Z*_*n,m*_does not include the sites at which both cells carry the same mutation, which occurs with probability of 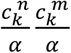.

As an index of the genetic variation within an individual tree, we then adopt the mean value of the pairwise genetic difference. The index evaluates the variability among offspring produced from the tree (i.e., seeds and young trees). The mean pairwise genetic difference is given as follows:

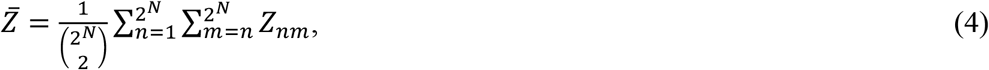

where 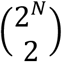is the number of possible pairs of branches within a tree having 2^*N*^ branches. We note that 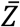 emphasizes mutations occurred in paths in the central portion than those in the marginal portion of the tree (Iwasa et al., 2023b).

#### (5) Pairwise genetic difference for a single stem cell case

We first consider a SAM consisting of a single stem cell (*α* = 1). For this case, all mutations accumulating in the stem cell are fixed in a SAM, resulting in genetic homogeneity within the SAM. The pairwise genetic difference between two SAMs thus consists of mutations that occurred during elongation after the forking of two branches. These elongation processes are on the shortest path connecting the SAMs of focal branches along the trunk and branches (Fig. 2A). We call this as “the path after forking” and its length is denoted by 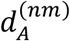, the physical distance between branch *n* and *m*. Based on Eq. (2), the number of genetic differences between branches *n* and *m, Z*_*n,m*_, is expressed as follows:

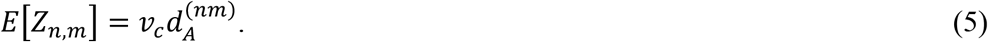

The expected number of genetic differences is proportional to the length of the path connecting the SAMs of the two focal branches. When plotted on a plane with the horizontal axis 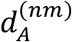, the points lie on a straight line passing through the origin.

**Figure 2.**
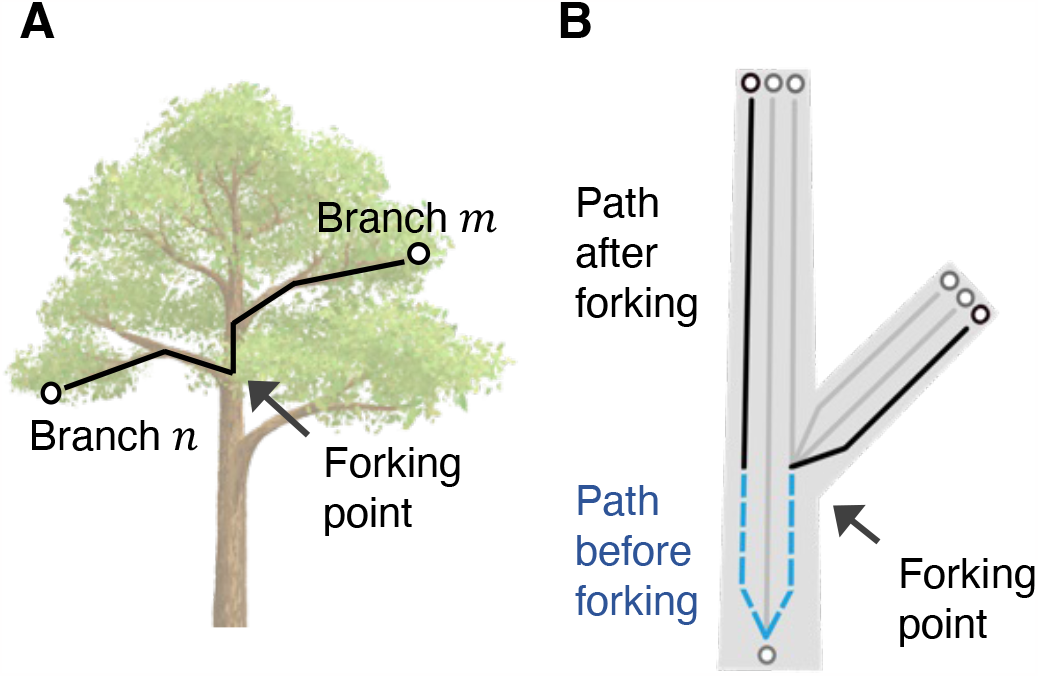
Schematic representation of paths for somatic mutation accumulation. (A) The shortest path connecting the SAMs of focal branches, shown in a black line, is called the path after forking; (B) The paths for the multiple stem cells case. Sampled stem cells and their paths are shown in black as circles and lines, respectively. Nonsampled cells and their paths are shown in grey. The path between apical and axillary meristems is illustrated. Sampling different stem cell lineages in each branch causes the contribution of a path before forking, shown in the blue dashed line.

#### (6) Effect of genetic heterogeneity among stem cells in the same SAM

We then consider a SAM having multiple stem cells (*α* > 1). Under the structured meristem, stem cell lineages are retained separately within a SAM and accumulate mutations independently, leading to genetic heterogeneity within a SAM. In such a case, mutations that occurred before forking also contributes to the genetic difference, and the pairwise genetic difference becomes higher than the homogenous case. The contribution of these mutations is expressed as an additional term in Eq. (5), as follows:

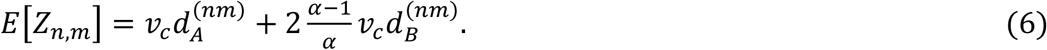

If we plot *E*[*Z*_*n,m*_ J against 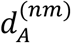, the term appears as the intercept of a linear line. We call elongation processes during which these mutations occur as the “path before forking.” This path, denoted as 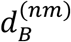, spans from the forking point to the common ancestral cell of sampled stem cells from each branch tip (Fig. 2B). Here, (*α* − 1)/*α* is the probability that sampled cells have their common ancestral cell before the forking point, and 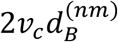 is the expected genetic difference occurred during the path before forking (see Appendix C in the Supplemental Material for derivation). If a SAM is genetically homogenous (i.e., *α* = 1), the term of the path before forking disappears, and Eq. (6) simplifies to Eq. (5). We calculated the pairwise genetic difference for all branch pairs within a single tree, based on Eq. (6), and evaluated the mean of them under varying branching architecture, based on Eq. (4) (for details see Appendices C and D in the Supplemental Material).

## RESULTS

### Pairwise genetic differences under different branching architectures

Figure 3 illustrates how the pairwise genetic differences are affected by varying branching architecture and the number of stem cells. Each part of Fig. 3 illustrates the result for all pair of branches in a single individual. The vertical axis indicates the pairwise genetic difference between branch tips *Z*_*n,m*_. The horizontal axis indicates the physical distance between branch tips, or the length of the path after forking 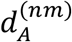. The values of *Z*_*n,m*_ depends on pair location within branching architecture. Pairs including the trunk tip are shown in triangles, and other pairs are shown in solid circles. To examine the effect of modifying the parameters controlling the architecture, we changed only one parameter with the others fixed and plotted the relationship in different parts of Fig. 3. Panels on the upper row (Fig. 3A, B) are for the single stem cell case (*α* = 1), and those on the lower row (Fig. 3C, D) are for the multiple stem cells case (*α* > 1).

**Figure 3.**
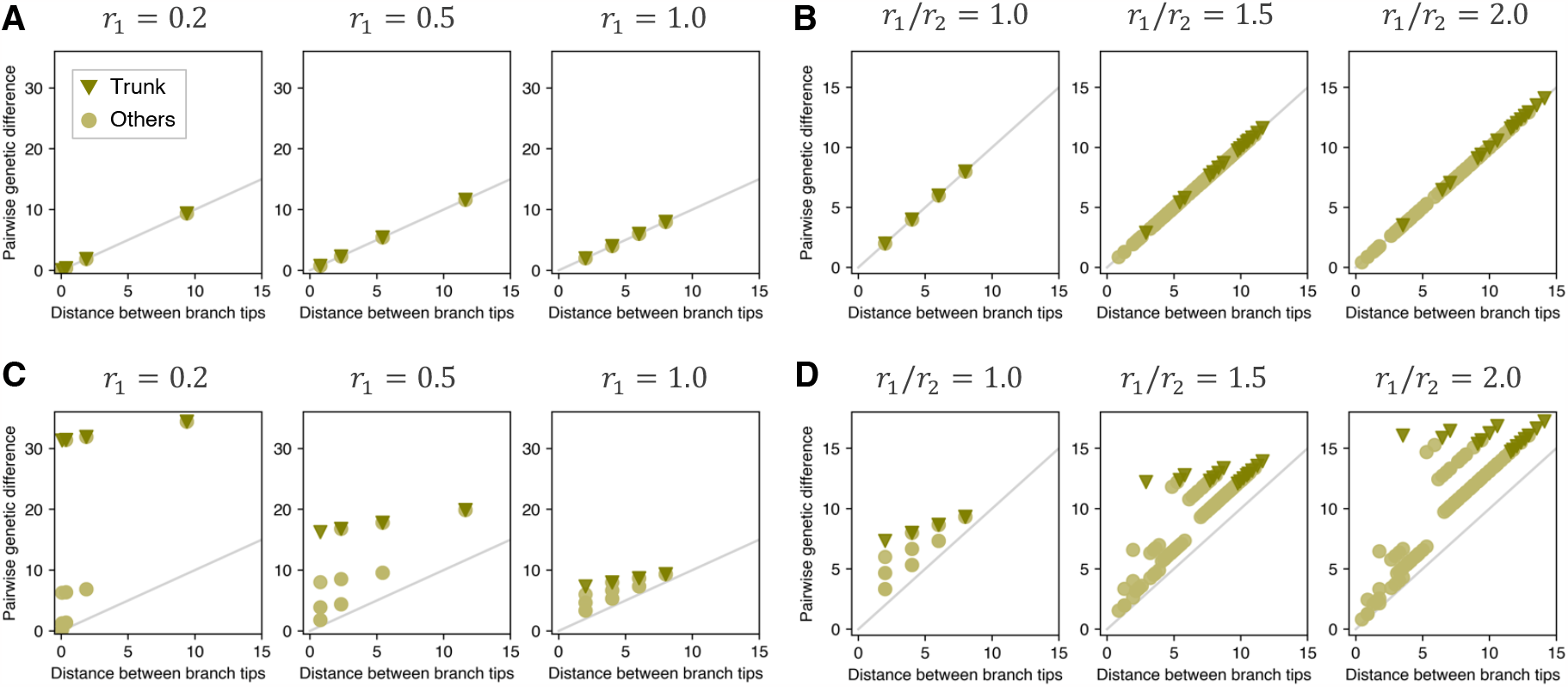
Relationship between pairwise generic difference and distance between branch tips, for the single stem cell case (*α* = 1; A, B) and the multiple stem cells case (*α* > 1; C, D). Vertical axis is the pairwise genetic difference, and horizontal axis is the physical distance between branch tips (the length of path after forking). Branch pairs including the trunk tip are denoted in triangle points, and other lateral branch pairs are denoted in circle point. The line is given in Eq. (5). (A) Pairwise genetic differences under different daughter-mother ratios, with *α* = 1 and *r*_1_/*r*_2_= 1; (B) Pairwise genetic differences under different main-lateral ratios, with *α* = 1 and *r*_1_= 1. (C) Pairwise genetic differences under different daughter-mother ratios, with *α* = 3 and *r*_1_/*r*_2_= 1; (D) Pairwise genetic differences under different main-lateral ratios, with *α* = 3 and *r*_1_= 1. Other parameters were *v*_*c*_ = 1 and *N* = 4.

#### (1) Single stem cell

For the single stem cell case, where a SAM is genetically homogenous, the pairwise genetic differences were proportional to the path after forking, as indicated in Eq. (5). As shown in Figs. 3A and 3B, the pairwise genetic differences lay on the straight line passing through the origin. Different branching architectures resulted in varying lengths of the path after forking, plotted in different values of the horizontal axis in Fig. 3.

Figure 3A illustrates the result when we modified the daughter-mother ratio *r*_1_ with others fixed. As DM ratio *r*_1_ increased, the differences between the most distant branches (points in the upper right) increased and then decreased. In contrast, the pairwise genetic differences between nearby branches monotonically increased (Fig. 3A).

Figure 3B illustrates the result when we modified the main-lateral ratio *r*_1_/*r*_2_, with others fixed. As ML ratio *r*_1_/*r*_2_ increased, the pairwise genetic differences moved to both positive and negative directions depending on the relative position of the sampled pairs. The genetic differences increased for pairs connected by the path including the main branch, especially for pairs including the trunk tip shown in triangles. However, the values decreased for the pairs between marginal lateral branches as their connected path do not include a portion of the main branch (Fig. 3B).

#### (2) Multiple stem cells

Figures 3C and 3D illustrate the results for the multiple stem cells case (*α* > 1), where a SAM is genetically heterogeneous. The straight line indicates the value given by Eq. (5), which is the same one for *α* = 1. The pairwise genetic differences were all above the straight line, as shown in Eq. (6). Changing parameters modified branching architectures and resulted in varying pairwise genetic differences. We again changed only one parameter with others fixed.

Compared with the case for *α* = 1, the pairwise genetic differences became larger for *α* > 1, and these differences were pronounced for smaller DM ratio *r*_1_ and larger ML ratio *r*_1_/*r*_2_ (Fig. 3C, D). For both ratios, the genetic differences between pairs of neighboring branches, which are close to each other, became higher, especially for the main branch that has a long path to the base of the tree.

### Within-individual genetic variation under different branching architectures

We evaluated the magnitude of genetic variation within an individual tree by 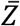, the mean value of the pairwise genetic difference *Z*_*n,m*_ for all pairs of branches within a tree, given in Eq. (4). The results are shown in Figs. 4A and 4B as the heat maps of 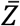, where two axes indicate parameters affecting branching architecture. Figures 4C and 4D illustrate how 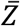 changed with a parameter on the horizontal axis with the other fixed.

**Figure 4.**
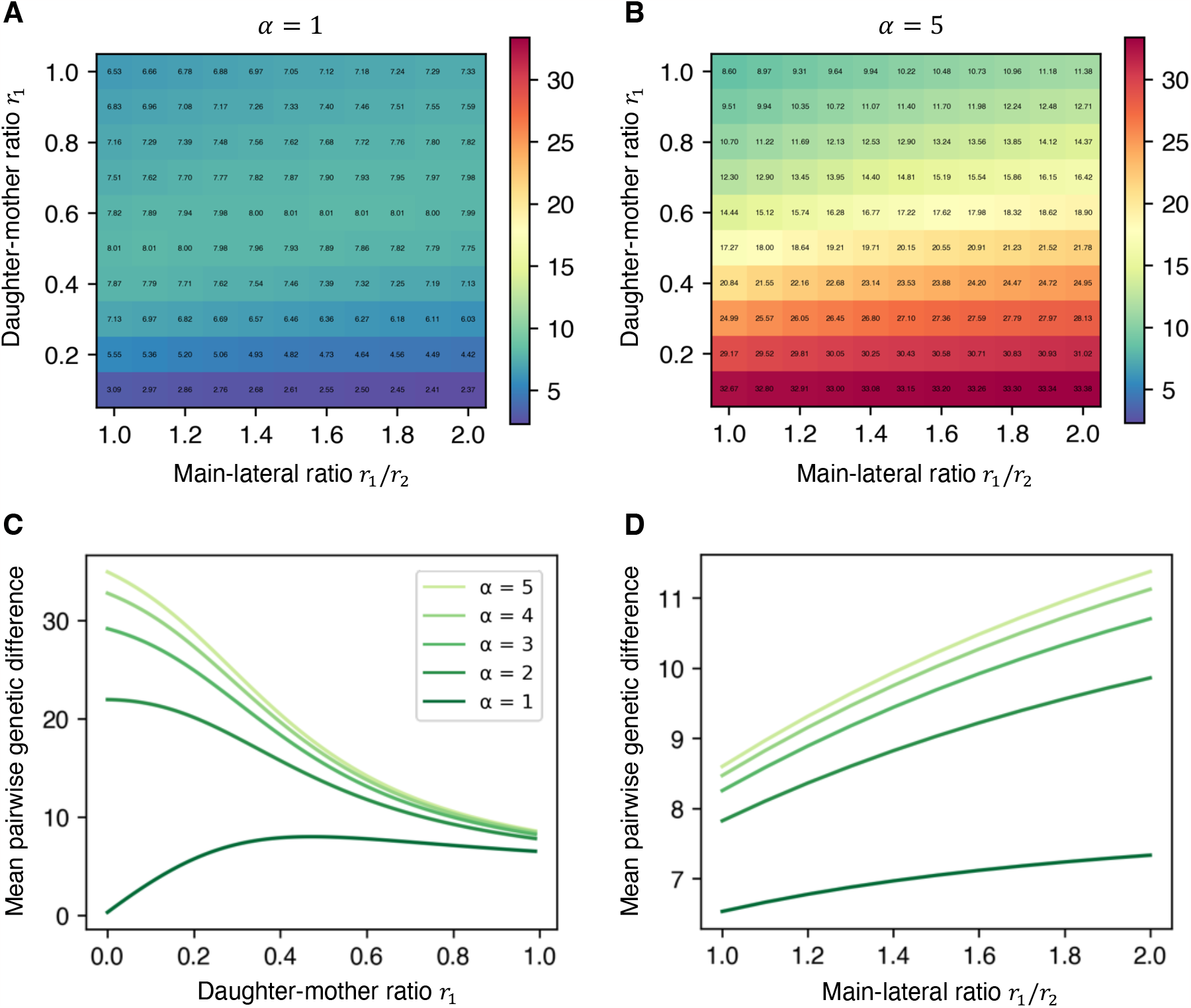
Genetic variation within an individual tree under different branching architectures. (A) Mean pairwise genetic difference for the single stem cell case (*α* = 1); (B) Mean pairwise genetic difference for the multiple stem cells case (*α* = 5); (C) Mean pairwise genetic difference with daughter-mother ratio *r*_1_, while keeping *r*_1_/*r*_2_ = 1; (D) Mean pairwise genetic difference with main-lateral ratio *r*_1_/*r*_2_, while keeping *r*_1_= 1. Lighter colored lines represent higher stem cell numbers. Other parameters were *v*_*c*_ = 1 and *N* = 4.

#### (1) Single stem cell

Figure 4A illustrates the effects of changing branching architectures on the mean pairwise genetic difference 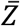 for the single stem cell case (*α* = 1). The vertical axis represents the daughter-mother ratio, and the horizontal axis represents the main-lateral ratio.

The DM ratio *r*_1_ influenced 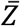 more strongly than the ML ratio *r*_1_/*r*_2_. Interestingly, the mean pairwise genetic difference 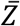 achieved the maximum at an intermediate value of the DM ratio. Additionally, 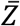 was close to zero for a very small DM ratio, whereas 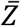 converged to a positive value for the larger DM ratio, as illustrated in the curve of *α* = 1 in Fig. 4C. These numerical results are confirmed by mathematical analysis in Appendix E in the SM. The value of *r*_1_ that attained the maximum 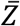 became larger as *r*_1_/*r*_2_ increased (Fig. 4A).

On the other hand, 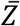 monotonically increased with the ML ratio *r*_1_/*r*_2_, as illustrated by the curve of *α* = 1 in Fig. 4D, although for very small DM ratios, 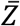 monotonically decreased with the ML ratio.

#### (2) Multiple stem cells

For the multiple stem cell case (*α* > 1), the path before forking contributes to 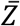 in addition to the path after forking. Figure 4B illustrates the effects of changing branching architectures on 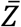 in the heatmap.

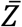 monotonically decreased with the DM ratio *r*_1_ and increased with the ML ratio *r*_1_/*r*_2_ (Fig. 4B), but it depended much more strongly on *r*_1_ than on *r*_1_/*r*_2_. The monotonic dependence on *r*_1_ for the multiple stem cell case (*α* > 1) was quite different from the single stem cell case (*α* = 1) discussed in the last subsection: 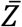 became much higher for small *r*_1_. These numerical results are confirmed by mathematical analysis in Appendix E in the SM. The dependence of each parameter is shown clearly in Figs. 4C and 4D.

## DISCUSSION

In this paper, we studied the magnitude of within-individual genetic variation accumulated in trees with different branching architectures. To describe the tree shape, we adopted a simple tree-like body model (Honda, 1971; Honda and Fisher, 1979), considering two parameters: daughter-mother ratio and main-lateral ratio. For different branching architectures, we kept the total length of branches constant. Focusing on two contrasting types of SAM with single and multiple stem cells, we evaluated the genetic variation of the whole tree by the mean pairwise genetic distance 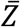. We note that this index tends to emphasize mutations occurred along paths in central portions than those in the marginal portions of a phylogenetic tree (Iwasa et al., 2023b).

### Effects of daughter-mother ratio and main-lateral ratio

One of the most interesting results we observed was that, for the single stem cell case (*α* = 1), the within-individual genetic variation 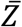 achieves the maximum for the intermediate value of daughter-mother ratio *r*_1_. Additionally, 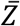 is lower for a very small *r*_1_ than that for a very large *r*_1_. If *r*_1_ is very small and close to zero, most mutations occur along the long base trunk, denoted by h, not along the subsequent daughter branches. Because mutations occurring in the base trunk do not contribute to 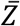 under *α* = 1, 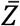 declines for very small *r*_1_ (Fig. 4A). In contrast, if *r*_1_ is large and close to one, mutations occur equally in all the branches. Most mutations are distributed to the terminal portions of branches, because descendent daughter branches are more numerous than the mother branches. However, those mutations are only shared with the small number of branch pairs and do not contribute the mean value of the pairwise genetic difference much. Thus, 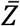 declines for larger *r*_1_ close to one. Consequently, the maximum value of 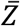 is achieved by an intermediate value of daughter-mother ratio.

The results are quite different for the multiple stem cells case (*α* > 1). Figure 4C indicates that 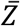 monotonically decreases with *r*_1_. Smaller *r*_1_ results in the longer base trunk (Fig. 1C), and under the heterogeneous SAM, mutations occurring in such a long portion contribute to the pairwise genetic difference as the path before forking. Consequently, 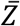 becomes higher for smaller *r*_1_, contrasting with low 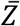 for the single stem cell case (*α* = 1) where only the path after forking contributes.

The second index, main-lateral ratio *r*_1_/*r*_2_, determines the degree of apical dominance of the branching architecture. As the ratio becomes larger, the tree becomes taller (Fig. 1D), and the within-individual genetic variation 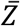 becomes larger. This trend is attributed, in part, to the increasing distances between main side branches, which compensate for the decreasing distances between lateral side branches (Fig. 4A). The increase of 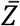 is more pronounced for *α* > 1, because the contribution of the path before forking increases due to the longer main branches (Fig. 1D).

### Genetically heterogenous SAM and path before forking

Because the number of stem cells is normally larger than one in trees (Dermen, 1969), the result of *α* > 1 appears to be more realistic than the result of *α* = 1. If so, the analysis concludes that the within-individual genetic variation should decrease with the DM ratio and increase with the ML ratio. However, we note that these results were obtained under the assumption that the stem cell lineages in a SAM remain separated from each other and never undergo replacement during elongation, and different stem cells in a SAM share a common ancestor only at the base of the branches (e.g., the base of the whole tree), as a result of successful asymmetric cell division of the structured meristem.

Sometimes, however, the stem cells may fail to leave their own successors. In such a case, one of the other stem cells duplicates, and the stem cell lineage is superseded by another stem cell lineage (Stewart and Dermen, 1970; Ruth et al., 1985; Rogers and Bonnett, 1989; Burian et al., 2016), decreasing the amount of genetic heterogeneity within the SAM. The common ancestor of two stem cells at the SAM can be located anywhere between the branch tip and the base of the tree. Iwasa et al. (2023) called the distance between the common ancestral cell and the sampled location as “coalescent length” and analyzed it mathematically. The mean coalescent length between two different stem cells converges to an equilibrium value that depends on the probability of a stem cell failing to leave behind its successor, denoted as *q*. A high failure rate leads to somatic genetic drift during elongation, resulting in the quick loss of genetic variation among stem cells within a SAM (i.e., stochastic meristem). Under such genetically homogenous a SAM, the tree behaves similarly to the case with *α* = 1 in the current study. In contrast, a lower failure rate maintains genetic heterogeneity within a SAM, resulting in behavior similar to the case with *α* > 1.

The results for a SAM having genetic heterogeneity (*α* > 1) indicate that mutations occurring in the path before forking enhance within-individual genetic variation, even when a tree has small expansion of branches (e.g., small *r*_1_ or large *r*_1_/*r*_2_). In dense forests with closed canopies, trees have limited expansion of lateral branches while the trunk elongates (Fisher and Hibbs, 1982). In trees with such an elongated shape, the contribution of paths before forking may lead to a high genetic variation within an individual tree, particularly when the tree retains genetic heterogeneity within a SAM. The importance of the path before forking is significant for trees living in dense closed-canopy forests, such as tropical forests. Whereas most previous empirical studies of somatic mutations have been performed on single-standing iconic trees in temperate zones, we need more empirical study of the within-individual genetic variation of trees in the dense forest.

### Phylogeny of somatic mutations

The phylogeny within a single tree reflects the accumulation pattern of somatic mutations. Our model can generate the phylogeny based on the matrix of genetic distance (see Eq. (D1) in Appendix D in the Supplemental Material). For the single stem cell case (*α* = 1), the expected shape of the phylogeny exactly mirrors the physical shape of the tree, as the genetic difference between branch tips is proportional to the physical distance (Fig. 3A, B). In contrast, our model for the multiple stem cells case (*α* > 1) shows that the phylogeny deviates from the physical shape due to the contribution of the path before forking (Fig. 3C, D). Whether the phylogeny reflects the physical shape depends on the degree of genetic heterogeneity among stem cells in a SAM. Conversely, the shape of a phylogeny may enable to infer the dynamics of stem cell lineages that determines the heterogeneity within a SAM. Empirical studies have, in fact, reported both concordance and discrepancies between phylogenetic and physical shapes (Zahrdadníková et al., 2020; Perez-Roman et al., 2021; Satake et al. 2023).

Moreover, our model for a SAM having genetic heterogeneity (*α* > 1) provides two important predictions. First, the genetic distance between neighboring branches is expected to be much longer than predicted by physical distance, as Figure 3 indicated that genetic differences between neighboring branches are above the regression line through the origin. Second, this lengthening effect is more pronounced for main branch pairs than for terminal branch pairs, due to the longer path before forking in the main branch. Interestingly, some empirical results corresponding to Fig. 3 are in line with our first prediction (see Supplementary Figure 4.1 in Orr et al., 2020; Fig. 2 in Satake et al. (2023)), although more detailed comparison is needed. Notably, our model can also be applicable to other modular organisms, including clonal plants (Schmid, 1990; Reusch et al., 2021), for inferring the dynamics of stem cell lineages.

### Limitation of the model studied in this paper

Our model neglects the occurrence of *de novo* mutations during branching, notwithstanding that an axillary meristem experiences some cell divisions during its formation (Burian et al., 2016). To describe tree species that accumulate as many mutations during branching as during elongation, branching should also contribute to mutation accumulation (e.g., Klekowski et al., 1989; Tomimoto and Satake, 2023).

Additionally, we did not consider the effects of the branch angle. However, a divergence angle of lateral branches, denoted as *α* in Honda (1971), can have significant impacts on the within-individual genetic variation, because the angle may determine the genetic similarity between stem cells which form lateral branches (Iwasa et al., 2023a).

The number of seeds produced by each branch may differ depending on the location of a branch, influencing the genetic composition of offspring generated from a single tree. To incorporate the variance of seed production between branches, we can extend our model by considering weights in each branch in Eq. (4). Additionally, we may discuss alternative indexes for the within-individual genetic variation. These are topics in a twin paper (Iwasa et al., 2023b).

## CONCLUSIONS

Our study demonstrated that branching architecture affects the amount of genetic diversity stored in an individual tree. Moreover, the effects of branching architecture depend strongly on the genetic heterogeneity among stem cells in a SAM, which enhances within-individual genetic variation via path before forking. Evaluation of how branching architecture affects the amount of standing genetic variation of a tree population will be worthy of study.

## Acknowledgements

This work was supported by JSPS KAKENHI Grant Number JP23KJ1722 to S.T., and Grant Number JP23H04966 to A. S. We thank the following people for their very helpful comments: R. Hayashi, C. Myotoishi, R. Imai, E. Sasaki., and K. Noshita.

## Author Contributions

**S. T**. Conceptualization, Methodology, Visualization, Writing – original draft, Funding acquisition; **Y. I**. Writing – review & editing, Supervision; **A.S**. Writing – review & editing, Supervision, Funding acquisition.

## Data Availability Statement

Because the research is entirely theoretical, no data is available. Mathematical analyses are explained in the Supplemental Material.

## Online Supporting Information

Additional supporting information may be found online in the Supporting Information section at the end of the article, including **Appendices A** (‘Adjusting the base length to keep the total length of branches constant’), **B** (‘Mutation accumulation in a single cell lineage during elongation’), **C** (‘Pairwise genetic difference between axillary meristems’), **D** (‘Genetic distances within a branching architecture’), and **E** (‘Mathematical analysis of the pairwise genetic difference’).

## Supplemental material for

## Appendix A Adjusting the base length to keep the total length of branches constant

To compare the different branching architectures, the total length of branches is kept constant by adjusting the base length h, the branch length from the base to the first branching (Fig. 1B). First, the sum of the total length is given as follows:

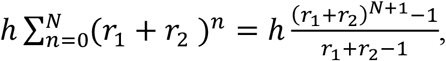

if *r*_1_+ *r*_2_ ≠ 1. Then, let h_8_ denote the base length of the branching architecture with *r*_1_= *r*_2_= 1, the total length of this architecture is h_0_(2^*N*+1^ − 1). This total length is used as a constant value. Namely, the base length h for each branching architecture is calculated to satisfy the following equation while varying *r*_1_/*r*_2_:

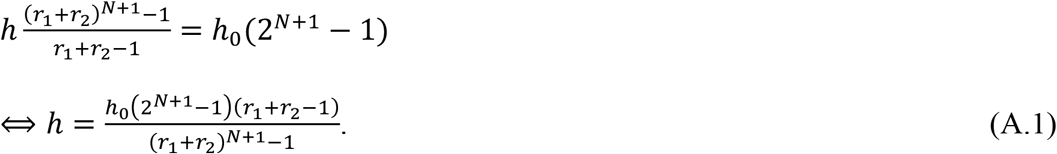

If *r*_1_ + *r*_2_ = 1, the sum of the total length is h(*N* + 1) and the base length is given as follows:

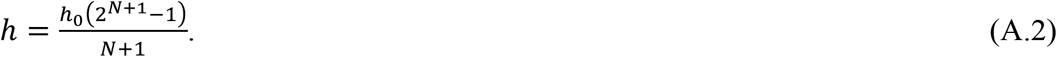

For simplicity, we set h_0_ = 1 in calculations. Based on these equations, we adjusted the base length h of different branching architectures in *r*_1_ / *r*_2_ and *r*_1_.

## Appendix B Mutation accumulation in a single cell lineage during elongation

During the process of elongation, somatic mutations occur and accumulate in each of the stem cell lineage. Consider the process of mutation accumulation in a single-cell lineage, which is a minimum unit of the mutation dynamics. Given that somatic mutations occur at a rate *μ*_*s*_ per site per cell division and the rate is small enough, the probability of a focal site being mutated after *t* cell divisions, *P*_*s*_(*t*), is approximated as follows:

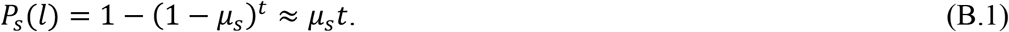

Then, the probability that *k* sites within the genome with size *G* are mutated, *P*_*G*_(*t*), follows the binomial distribution:

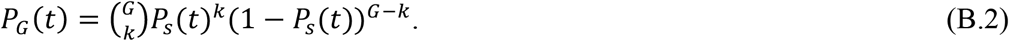

The expected number of somatic mutations accumulated in a single cell lineage after *t* cell divisions, *M*(*t*), is given as *M*(*t*) = *GP*_*s*_(*t*). By introducing the mutation rate per cell *μ*_+_ that equals to *Gμ*_*s*_, *M*(*t*) is given as follows:

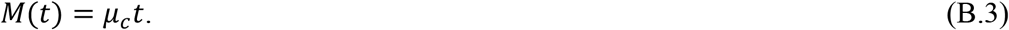

Given *η* is the stem cell division rate per unit branch growth, a stem cell divides *ηd* times for the branch to elongate distance *d*. Hence, the expected number of somatic mutations after *d* distance growth is given as *M*(*ηd*) = *μ*_*c*_*ηd*. By replacing *μ*_*c*_*η* to *v*_*c*_, the mutation rate per cell per unit distance growth, the expected number of somatic mutations after *d* distance growth is given as follows:

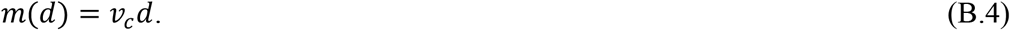

We clarify that our model addresses mutation accumulation linked to cell divisions. Regarding time-dependent mutations discussed in Satake et al. (2023), which accumulate independent of cell divisions, we assume uniform time requirements for unit growth across branches and different individuals with varying architectures.

## Appendix C Pairwise genetic difference for multiple stem cells case

When a SAM has multiple stem cells, the axillary meristem is generated from a subset of stem cell lineages in the apical meristem. Then, the axillary meristem does not inherit mutations accumulated in the other lineages that were not involved in the meristem formation. These mutations can contribute to the genetic difference between the apical and axillary meristems, because the mutations in the apical are not in the axillary meristem. Hence, even just after the forking, the apical meristem genetically differs from the axillary meristem without occurring *de novo* mutations. In Eq. (4), the contribution of those mutations is expressed as an additional term denoted by *B*_*f*_, as follows:

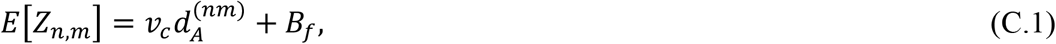

If we plot *E*[*Z*_*n,m*_J against 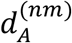, the term appears as the intercept of a regression line. We call elongation processes during which these mutations occur as the “path before forking” (Fig. 2B). In the following, we derive the explicit formula of *B*_*f*_.

To emphasize the genetic heterogeneity in the apical meristem, our model assumes that (i) all stem cell lineages are retained during elongation. Additionally, to simply describe the process of branching, our model assumes that (iii) a newly formed axillary meristem originates from a single stem cell in the apical meristem. Hence, only mutations accumulated in the single line are passed to and fixed in the axillary meristem, losing genetic heterogeneity.

Based on these assumptions, the path before forking spans from the forking point to a point of meristem formation (e.g., a branching point and the base of the tree), which corresponds to the common ancestral stem cell for both meristems (Fig. 2B). Let *d*_*B*_ be the distance of elongation from the point of meristem formation to the forking point. During *d*_*b*_ elongation prior to the forking event, each stem cells separately accumulates somatic mutations, increasing heterogeneity within the SAM. The stem cells are expected to genetically differ from each other by *v*_*c*_ (*d*_*B*_ + *d*_*B*_), based on Eq. (3).

Here, we consider the genetic difference between the apical meristem and the newly formed axillary meristem, just after the forking event. Because the axillary meristem originates from a single stem cell lineage in the apical meristem, the probability of sampling other cell lineages from the apical meristem is (*α* − 1)/*α*. Hence, the average heterozygosity between randomly sampled stem cells is calculated as 2*v*_*c*_*d*_*B*_ ×(*α* − 1)/*α* + 0 × 1/*α*, and the contribution of the path before forking is given as follows:

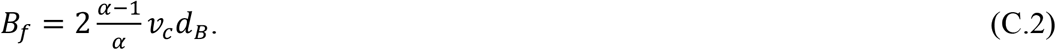

Does this Eq. (C.2) hold for the pairwise genetic difference not only between the apical and axillary meristems but also between axillary meristems? Figure C1 indicates the axillary meristems, both originating from a single apical meristem. If the axillary meristems are both formed from the same stem cell lineage with a probability of 1/*α*, a stem cell in each meristem shares the same mutations, and the path before forking dose not contribute to the pairwise genetic difference (Fig. C1A). However, if the axillary meristems are formed from different stem cell lineages with a probability of (*α* − 1)/*α*, a stem cell in each meristem differs by 2*v*_+_*d*_7_ mutations, and the path before forking contribute to the pairwise genetic difference (Fig. C1B). Thus, the expected contribution of the path before forking is 2*v*_*c*_*d*_*B*_ × (*α* − 1)/*α* + 0 × 1/*α*, yielding the same result as Eq. (C.2). Because the path after forking is also invariant even if focal branches are both axillary meristems, the expected pairwise genetic difference between SAMs at branches *n* and *m* is expressed as follows:

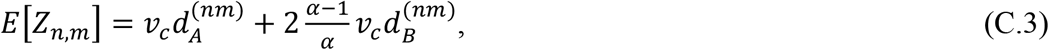

regardless of the relative position of the focal branches. Here, 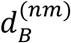 represents the path before forking between branches *n* and *m*.

Notably, our model for *α* > 1 assumes the type of branching called monopodial branching, which is a common way of branching in trees. In contrast, sympodial branching is a different type of branching where the axillary meristem replaces the apical meristem and produce a new main branch (Barthélémy and Caraglio, 2007). In tree species with sympodial branching, the genetic heterogeneity within the meristem may be lost during every step of elongation, even for the apical meristem (Zahrdadníková et al., 2020). Hence, the genetic difference within these trees can be modelled by the single stem cell case (*α* = 1).

**Figure C.1.**
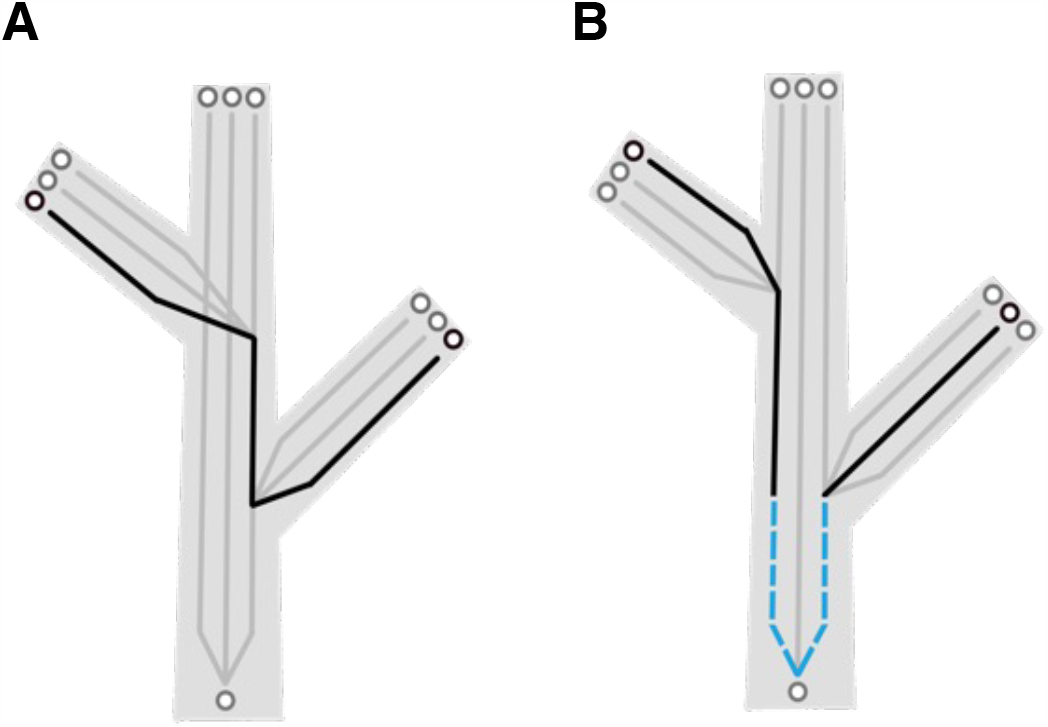
Schematic representation of paths between axillary meristems. Sampled stem cells and their paths are shown in black as circles and lines, respectively. Non-sampled cells and their paths are shown in grey. (A) The axillary meristems are formed from the same stem cell lineage resulting in no contribution of the path before forking; (B) The axillary meristems are formed from the different stem cell lineages. The path before forking is denoted in the blue dashed line.

## Appendix D Genetic distances within a branching architecture

The pairwise genetic difference is extended to all branches within an individual tree by the distance matrix, whose *nm* element is given in Eq. (6). Namely, the matrix of pairwise genetic difference is expressed as follows:

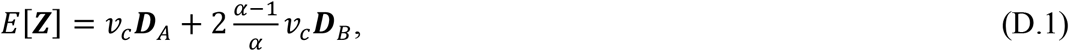

where 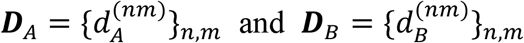 are the distance matrices for the path of after and before forking, respectively. The matrices ***D***_*A*_ and ***D***_*B*_ are calculated based on the model of branching architecture discussed above. As an example, we show a branching architecture with four branches, formed by two times reiteration of bifurcation process, *N* = 2 (Fig. 1B). The distance matrices for the path after forking is given as follows:

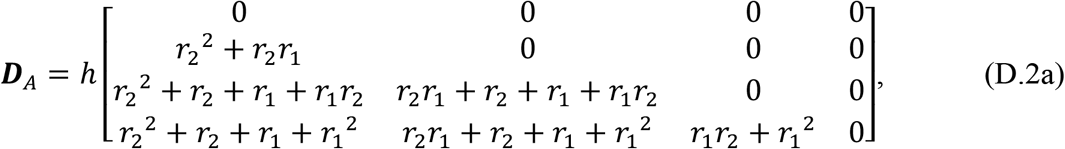

and the distance matrices for the path before forking is given as follows:

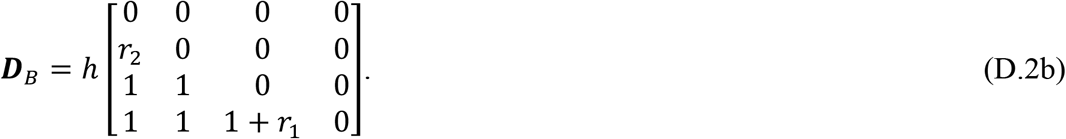

Here, h is given in Eq. (1) for certain values of *r*_1_ and *r*_2_. The order of the row and column of the matrices corresponds to the branch IDs in Fig. 1B. Notably, the matrix in Eq. (D.1) gives the expected shape of the phylogeny within a single tree, because the matrix shows the genetic distance between tips of branches.

## Appendix E Mathematical analysis of the pairwise genetic difference

Here, we verified the numerical results in the main manuscript by directly analyzing the equation for the pairwise genetic differences for *N* = 2, given in Eq. (D.1) and Eq. (D.2). Since,

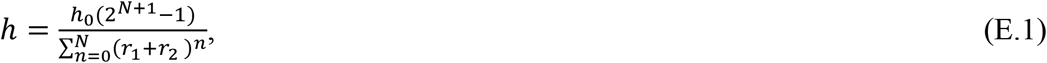

The pairwise genetic differences, given in Eq. (D.1) and Eq. (D.2), are expressed as the function of *r*_1_and *r*_2_as follows:

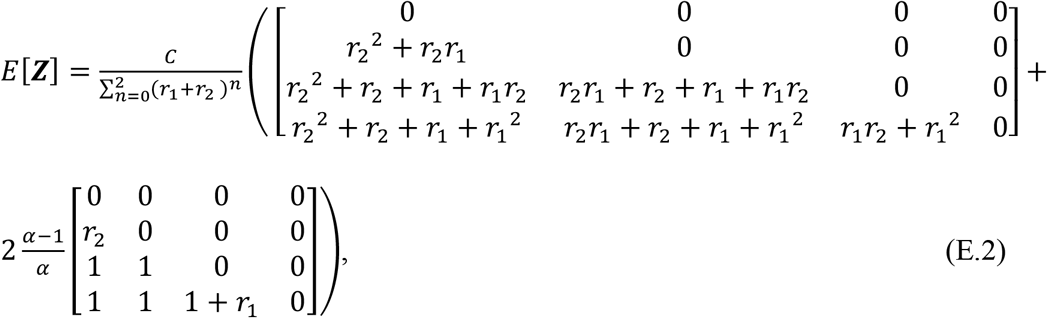

where *C* = *v*_*c*_h_0_(2^*N*+1^ − 1). Then, based on 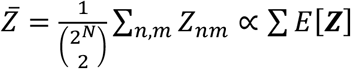, we first consider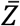 for the case where the daughter-mother ratio is close to zero (*r*_1_→ 0). Since *r*_1_≥ *r*_2_

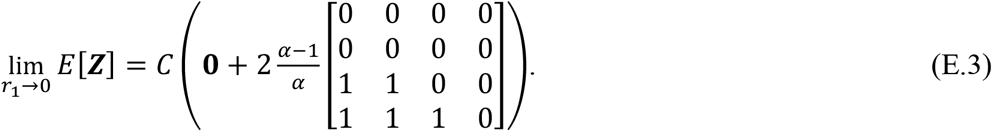

Then, 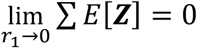, if *α* = 1, and 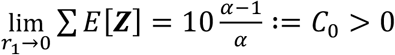, if *α* > 1.

This result clearly shows the effect of the path before forking for the multiple stem cells case, which are on the trunk (i.e., the portion of the base length; see Fig. (1B)). We then consider 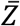 for the case where DM-ratio is close to one (*r*_1_ → 1). For any *r*_2_ ≤ *r*_2_,

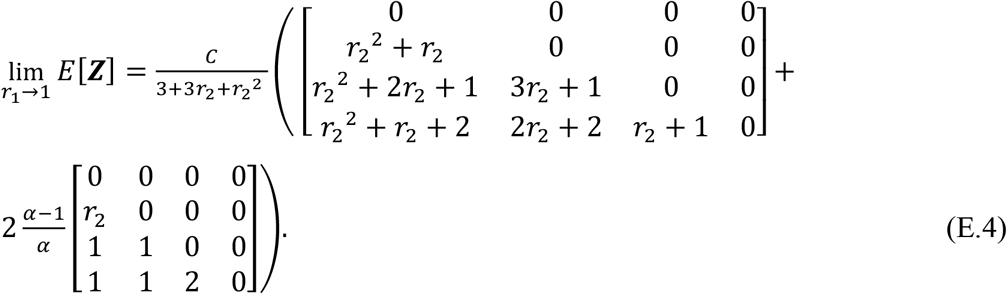

Thus, 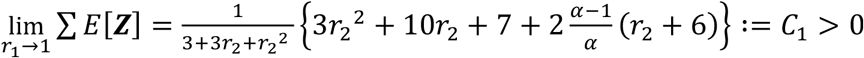, regardless of *α*. Additionally,

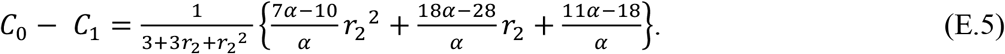

Therefore, if *α* = 1, *C*_0_ – *C*_1_ < 0; and if *α* > 1, *C*_0_ – *C*_1_ > 0.

To sum up, if 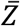 converges to zero for *r*_1_→ 0 and converges to the positive value for *r*_1_→ 1. In contrast, if 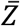converges to the positive value for *r*_1_ → 0 and converges to the positive value for *r*_1_→ 1.

We also verified the results by calculating derivatives. When *α* > 1, the derivative of Eq. (E.2) is negative for *r*_1_→ 0 and *r*_1_→ 1, indicating the monotonic decreasing 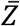 for *r*_1_. In contrast, when *α* = 1, the derivative of Eq. (E.2) is positive for *r*_1_→ 0, indicating the increasing trend of 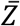for *r*_1_which is small enough. However, the derivative is also positive for *r*_1_ → 1. Hence, when *N* = 2 and *α* = 1, 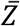 monotonically increases for *r*_1_ and does not have the maximum at an intermediate value of *r*_1_, as the *N* = 4 case in the numerical calculation. The derivative of Eq. (E.2) is expressed as follows:

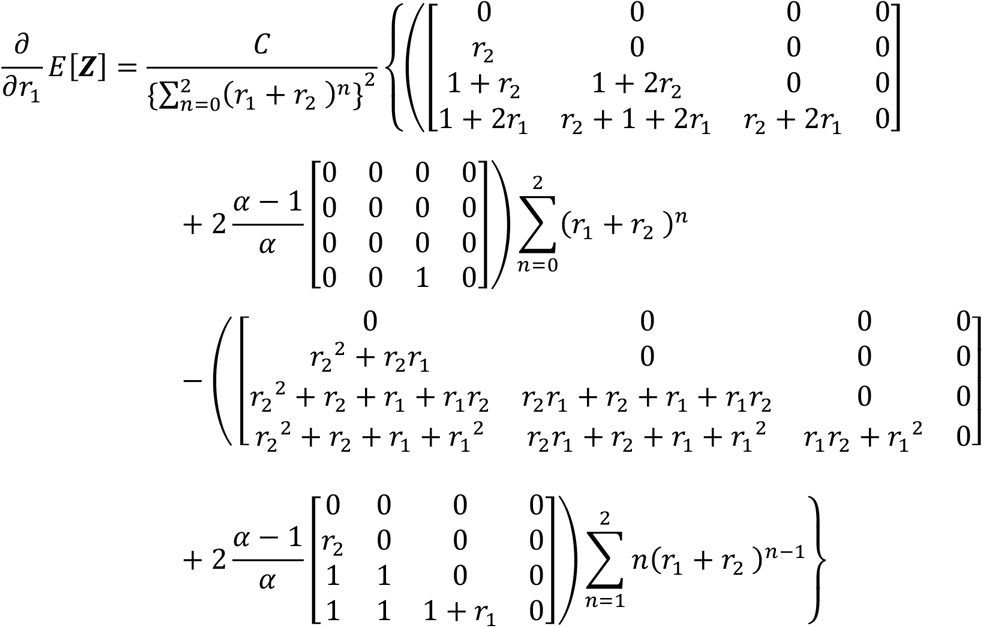

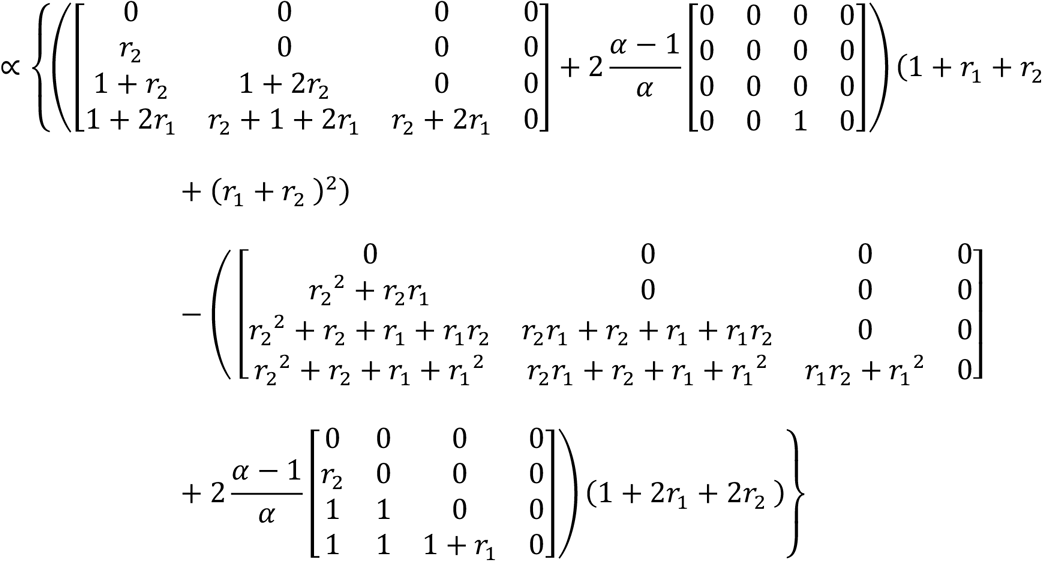

For small *r*_1_, since *r*_1_ ≥ *r*_2_,

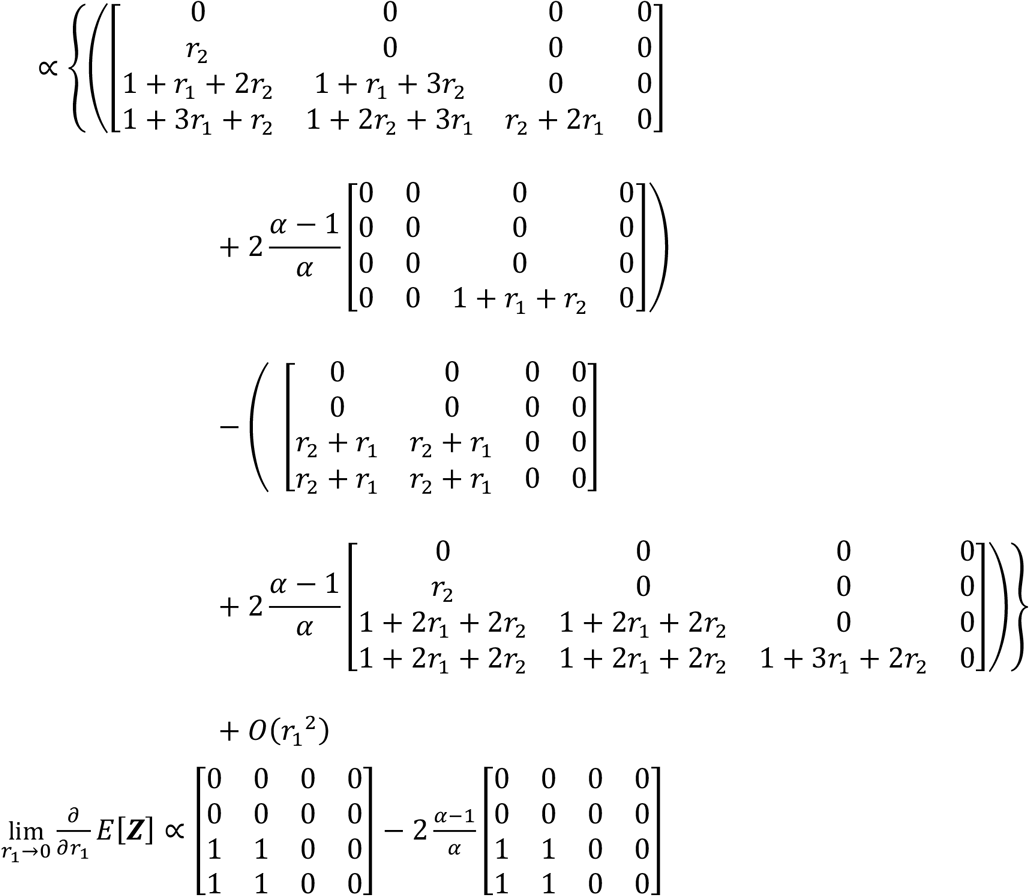

Hence, if 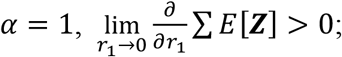 and if 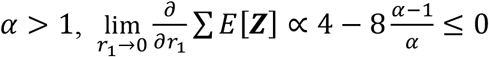 (the equality holds when *α* = 2). This result is in line with Fig. 4C in the manuscript. For *r*_1_, *r*_2_→ 1,

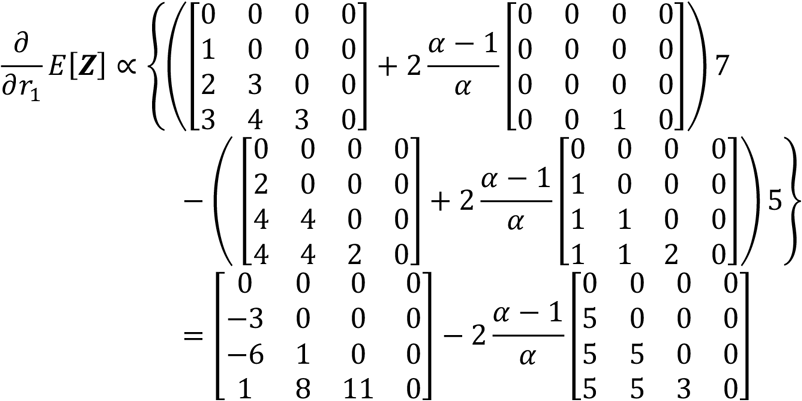

Hence, if 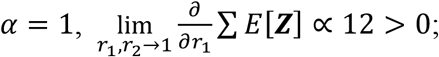 and if 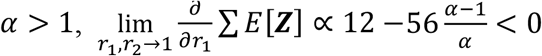.

## Notes

### Competing Interest Statement

The authors have declared no competing interest.

